# Antibody reliability influences observed mRNA-protein correlations in tumour samples

**DOI:** 10.1101/2022.12.23.521733

**Authors:** Swathi Ramachandra Upadhya, Colm J. Ryan

## Abstract

Reverse phase protein arrays (RPPA) have been used to quantify the abundances of hundreds of proteins across thousands of tumour samples in the Cancer Genome Atlas (TCGA). By number of samples, this is the largest tumour proteomic dataset available and it provides an opportunity to systematically assess the correlation between mRNA and protein abundances. However, the RPPA approach is highly dependent on antibody reliability and approximately one third of the antibodies used in the TCGA are deemed to be somewhat less reliable. Here, we assess the impact of antibody reliability on observed mRNA-protein correlations. We find that, in general, proteins measured with less reliable antibodies have lower observed mRNA-protein correlations. This is not true of the same proteins when measured using mass spectrometry. Furthermore, in cell lines, we find that when the same protein is quantified by both mass spectrometry and RPPA, the overall correlation between the two measurements is lower for proteins measured with less reliable antibodies. Overall our results reinforce the need for caution in using RPPA measurements from less reliable antibodies.

## Introduction

Proteins are the primary actors in our cells and their activities are often deregulated in cancer. Quantifying proteomic abundances in patient tumours can aid the identification of pathways and processes that are deregulated in specific tumours while, across cohorts of patients, variation in protein abundances can be linked with specific phenotypic outcomes and responses to therapy. Unfortunately, our ability to quantify protein abundances in large cohorts has lagged significantly behind our ability to quantify mRNA abundances. Consequently, mRNA abundances are frequently used as proxies for protein abundances. However, there is only a moderate correlation between the two measurements with this limited correlation attributed to a combination of post-transcriptional regulation and limitations in our ability to accurately and reproducibly quantify transcripts and proteins (Vogel & Marcotte, 2012; Liu *et al*, 2016; Franks *et al*, 2017; Buccitelli & Selbach, 2020; Csárdi *et al*, 2015; Upadhya & Ryan, 2022).

The two main approaches that have been employed to quantify protein abundances in patient cohorts are mass spectrometry (MS), which has been recently used to quantify the abundance of thousands of proteins in up to a couple of hundred patients (Ellis *et al*, 2013; Petralia *et al*, 2020), and reverse phase protein arrays (RPPA), which have been used to quantify the abundances of a couple of hundred proteins in thousands of patients (Li *et al*, 2013; Chen *et al*, 2019). In recent work we have shown how limitations in our ability to reproducibly quantify proteins using mass spectrometry contributes to the moderate observed correlation between mRNA and protein abundances (Upadhya & Ryan, 2022). Here we focus our analysis on technical factors associated with protein abundances measured using RPPA.

RPPA is a simple and cost effective antibody based quantification approach that can be used to profile large numbers of samples. The Cancer Genome Atlas (TCGA) program has used RPPA to quantify the abundances of ∼200 proteins for >7,500 tumour samples across 32 different cancer types (Chen *et al*, 2019). In addition to RPPA profiles, the vast majority of the TCGA samples have linked exome sequencing, copy number profiles and transcriptomes, facilitating integrated analyses of different molecular profiles (Cancer Genome Atlas Research Network, 2011; Cancer Genome Atlas Research Network *et al*, 2013a; Ge *et al*, 2018). Among other applications, the TCGA RPPA profiles have been used to classify tumours into cancer types (Zhang *et al*, 2015), to understand the differential activation of signalling pathways across tumours (Zhang *et al*, 2017), to identify mechanisms of epithelial-mesenchymal transition (Koplev *et al*, 2018), and to assess the correlation between mRNA and protein abundances (Chen *et al*, 2019). Although the number of proteins quantified in this resource is relatively small, the large number of samples with linked RPPA profiles and transcriptomes provides a unique resource for understanding how mRNA variation contributes to protein variation.

A key factor in the accuracy of RPPA profiles is the reliability of the antibodies used, as even minor non-specific binding of antibodies can distort the signal from the target protein (Mannsperger *et al*, 2010). Antibody quality is a challenge for many biological assays, including Western blotting, but the problem is more acute for RPPA studies (Mannsperger *et al*, 2010). In Western blots, non-specific binding of antibodies can be somewhat addressed by focusing on results corresponding to the expected molecular weight of the protein assayed, something which is not possible with RPPA. Furthermore, with many assays the incubation conditions can be optimised for individual antibodies but this is not possible with the hundreds of antibodies used in RPPA assays (Mannsperger *et al*, 2010; Akbani *et al*, 2014a).

Western blots are frequently used to validate antibody specificity – with more reliable antibodies producing a single defined band at the correct molecular weight for the target protein. RNA interference, or knockout cell lines, can also be used to ensure that the target protein’s abundance is reduced when its production is inhibited (Mannsperger *et al*, 2010; Bordeaux *et al*, 2010). Within the TCGA RPPA pipeline, antibodies are systematically assessed using Western blots to ensure that each antibody produces a single or dominant band. Furthermore, they are assessed across a panel of cell lines to ensure that RPPA measurements of protein abundances are highly correlated with the same proteins when measured by Western blots.

Here, to understand the influence of the reliability of antibodies on observed mRNA-protein correlations, we analyse studies containing mRNA and protein expression profiles from The Cancer Genome Pan-Cancer Atlas (TCGA Pan-Can) (Cancer Genome Atlas Research Network *et al*, 2013b), and the Clinical Proteomic Tumour Analysis Consortium (CPTAC) (Ellis *et al*, 2013). We find that proteins that are quantified using less reliable antibodies have lower mRNA-protein correlations when using RPPA measurements whereas no such trend can be observed for the same proteins when quantified using mass spectrometry. These proteins do not appear to be less reliably quantified by mass spectrometry nor are they overall less abundant. By analysing data in cancer cell lines we find that proteins measured using less reliable antibodies tend to have lower correlation with mass spectrometry measurements of the same proteins. Overall, our results are consistent with antibody reliability influencing the accuracy of protein abundance measurements and reinforces the need for caution when analysing RPPA measurements made with less reliable antibodies.

## Results

### About one third of the antibodies used to quantify proteins using RPPA are not reliable

Although collectively they are often referred to as ‘high quality antibodies’ (Akbani *et al*, 2014b; Zhang *et al*, 2017; Şenbabaoğlu *et al*, 2016; Cancer Genome Atlas Research Network, 2017), the quality of the antibodies used for TCGA RPPA studies vary. All antibodies used are systematically assessed (Li *et al*, 2013; Akbani *et al*, 2014b; Chen *et al*, 2019). The two minimum criteria for validating antibody specificity used by the TCGA are (i) single or dominant band in a Western blot around the expected molecular weight of the target protein and (ii) a good Pearson correlation (>0.7) between abundances measured by RPPA and Western blotting across multiple samples (Li *et al*, 2013; Akbani *et al*, 2014a).

Based on these criteria antibodies are either discarded as unfit for use, categorised as ‘Valid’, or categorised as ‘Use with Caution’. Although the performance of ‘Use with Caution’ antibodies is poorer than those categorised as ‘Valid’, they are still used for quantification, typically because they bind to a protein known to have an important role in cancer.

The number of antibodies quantified in TCGA RPPA studies varies across different cancer types. Here we focus on a set of 180 antibodies that are used to measure (phospho)proteins across all TCGA Pan-Cancer studies. We restricted our analyses to those antibodies that are annotated as measuring the abundance of a single, non-phosphorylated protein and for which a quality evaluation is available (See Materials and Methods section). Of the 114 antibodies in this category, are labelled as ‘Use with Caution’ while 0 are labelled as ‘Valid’, i.e. ∼ 0 of antibodies used should be considered as less reliable (Fig 1, Table S1). The antibodies labelled as ‘Use with Caution’ include those that are used to measure the protein products of the oncogenes *MYC* and *BRAF* as well as the tumour suppressors *BRCA2* and *VHL* (Table S1).

**Figure 1.**
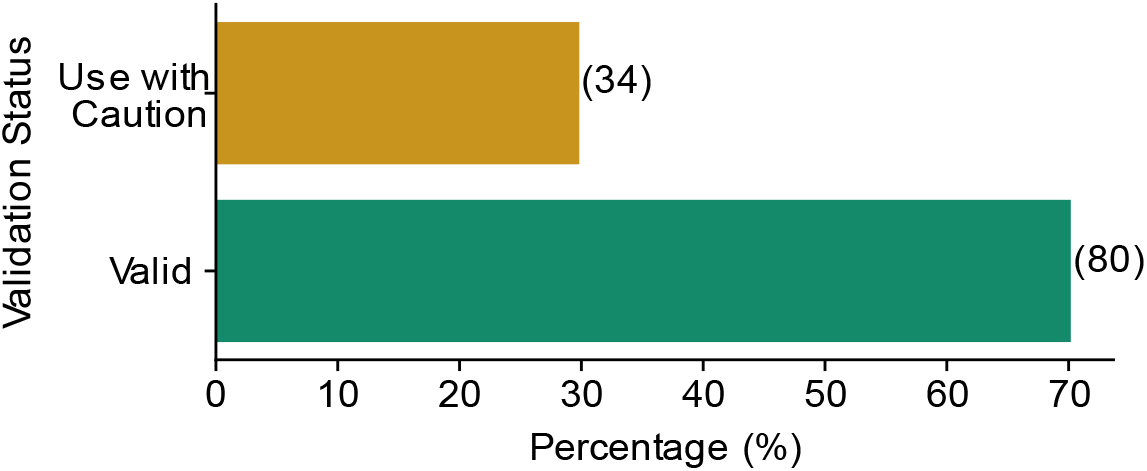
One third of antibodies used to quantify proteins in the TCGA RPPA Pan-Cancer dataset are not reliable. Bar chart showing the percentage of antibodies marked as ‘Valid’ and ‘Use with Caution’ for the TCGA Pan-Cancer studies. Only antibodies that are reported to target a single protein are included in this figure, those that target protein families (e.g. AKT1/2/3) and phosphoproteins are excluded. The actual number of antibodies in each class is shown in brackets.

### Proteins quantified using RPPA with valid antibodies have a higher mRNA-protein correlation

The reliability of the antibody used to quantify protein abundances will impact all downstream analyses of protein measurements including the analysis of mRNA-protein correlations. To assess the mRNA-protein correlation with respect to antibody validation status, we analysed the six TCGA Pan-Cancer Atlas studies with the highest number of samples profiled by RPPA – breast cancer (Cancer Genome Atlas Network, 2012b), ovarian cancer (Cancer Genome Atlas Research Network, 2011), colorectal adenocarcinoma (Cancer Genome Atlas Network, 2012a), endometrial carcinoma (Cancer Genome Atlas Research Network *et al*, 2013a), kidney renal cell carcinoma (Cancer Genome Atlas Research Network, 2013), and low grade glioma (Cancer Genome Atlas Research Network *et al*, 2015). We quantified the mRNA-protein correlation for all individual, non-phosphorylated proteins measured by RPPA and found that the median mRNA-protein correlation for proteins with ‘Valid’ antibodies was significantly higher compared to proteins with antibodies marked as ‘Use with Caution’ p-value ≤ 0.01 in all cases, two-sided Mann-Whitney U test) (Fig 2). We note that here we are referring to correlations calculated between the mRNA abundance and protein abundance of an individual protein across all samples within a study ‘across-sample’ correlation) (Vogel & Marcotte, 2012; Liu *et al*, 2016; Franks *et al*, 2017; Buccitelli & Selbach, 2020).

**Figure 2.**
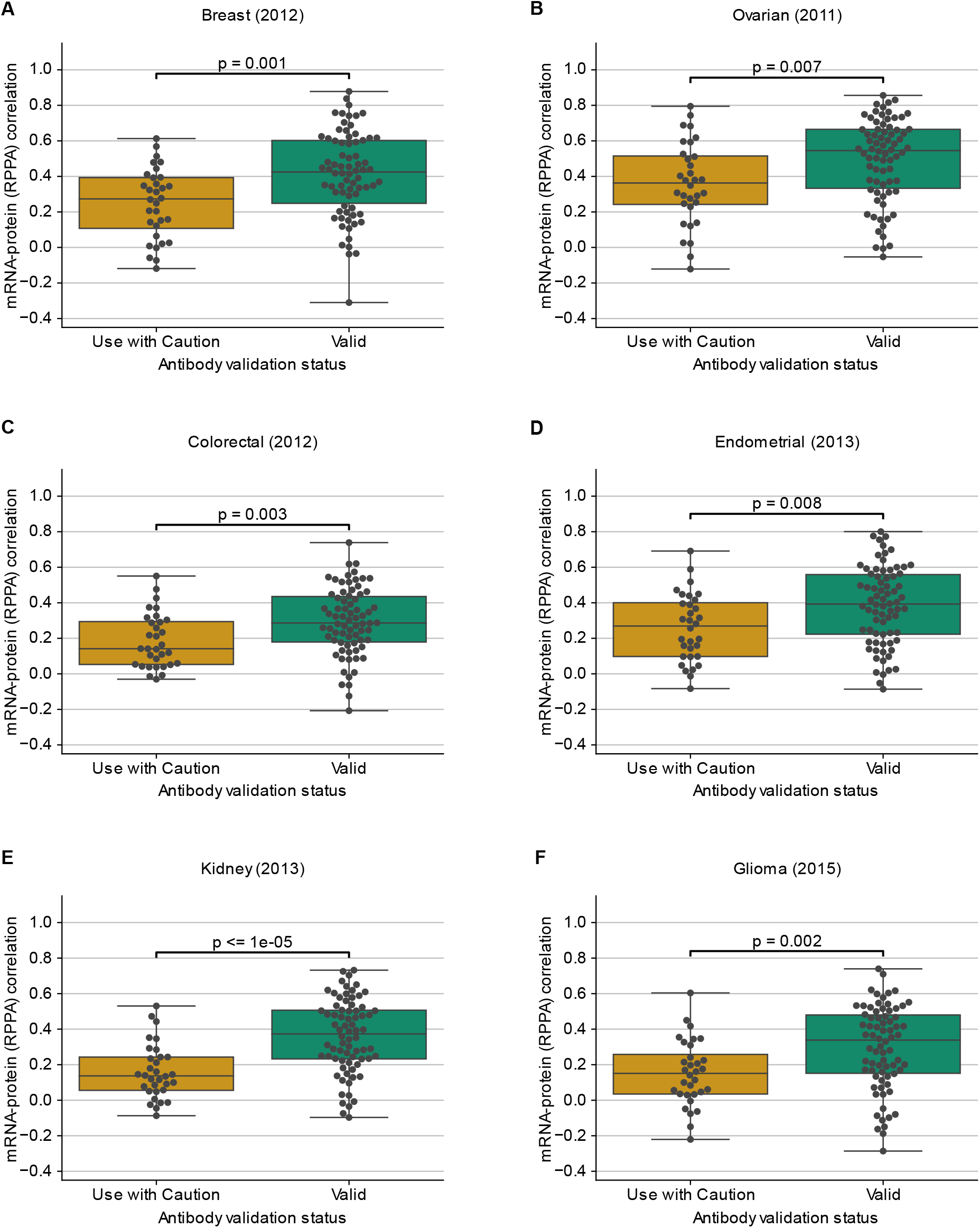
Proteins measured with less reliable antibodies have significantly lower mRNA-protein correlation. Boxplots showing the distribution of mRNA-protein correlations for proteins quantified using antibodies labelled as ‘Valid’ or ‘Use with Caution’. Correlation is measured using pearman’s correlation across all patients with measurements for both mRNA and protein in breast cancer (A), ovarian cancer (B), colorectal adenocarcinoma (C), endometrial carcinoma (D), kidney renal cell carcinoma (E) and low grade glioma (F). For each box plot, the black central line represents the median, the top and bottom lines represent the first and third quartiles, and the whiskers extend to 1.5 times the interquartile range past the box. Each point on a box plot represents a protein.

There are many factors, both technical and biological, that influence the variation in mRNA-protein correlations across proteins (Vogel & Marcotte, 2012; Liu *et al*, 2016; Buccitelli & Selbach, 2020; Franks *et al*, 2017; Upadhya & Ryan, 2022). However, we reasoned that if less reliable antibodies were the primary cause for the differences observed in Fig 2, then we should not see the same difference between the two groups of proteins when using protein abundance measurements quantified by mass spectrometry. To assess this, we analysed six CPTAC studies that either match the cancer type assayed in the TCGA RPPA studies (breast cancer (Mertins *et al*, 2016), ovarian cancer (Zhang *et al*, 2016), colorectal adenocarcinoma (Zhang *et al*, 2014), endometrial carcinoma (Dou *et al*, 2020), clear cell renal cell carcinoma (Clark *et al*, 2019)) or that assay a related cancer type (glioblastoma (Wang *et al*, 2021)). On repeating the above analysis on the mRNA-protein correlations for the CPTAC studies, we observed that the two classes of proteins had no particular pattern in the distribution of mRNA-protein correlation (Fig 3). In five out of six studies analysed there was no significant difference between the two groups, while in a single cancer type there was (p-value = 0.02, Mann-Whitney U test, two-sided).

**Figure 3.**
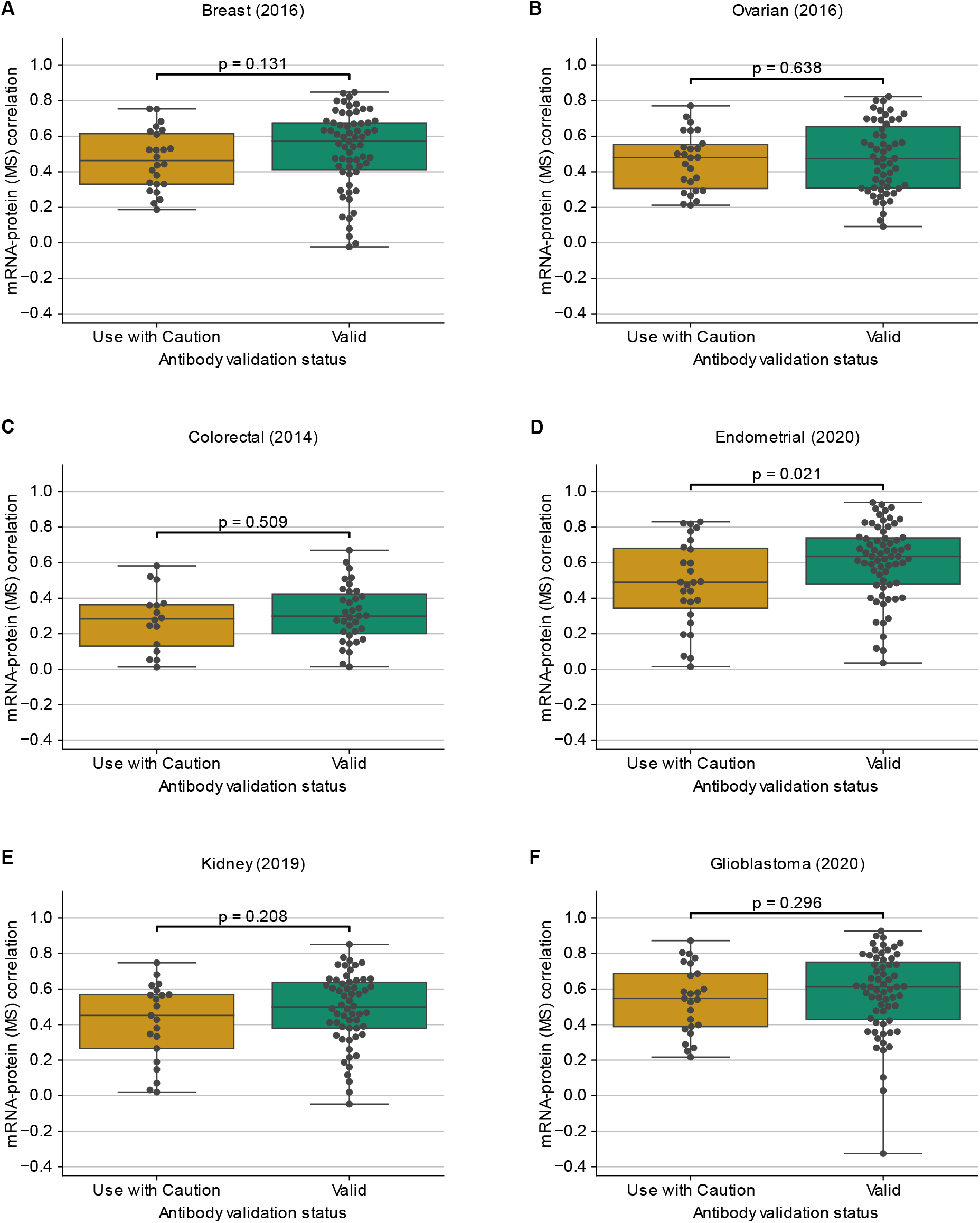
mRNA-protein correlations for proteins measured by mass spectrometry are not significantly influenced by the associated antibody validation status. Boxplots showing the distribution of mRNA-protein correlations wherein proteins are classified based on their antibody validation status and quantified using mass spectrometry. Correlation is measured using pearman’s correlation across all patients with measurements for both mRNA and protein in breast cancer (A), ovarian cancer (B), colorectal adenocarcinoma (C), endometrial carcinoma (D), clear cell renal cell carcinoma (E) and glioblastoma (F). For each box plot, the black central line represents the median, the top and bottom lines represent the first and third quartiles, and the whiskers extend to 1.5 times the interquartile range past the box. Each point on a box plot represents a protein.

The results in Fig 2 suggest that protein measurements made with antibodies marked as ‘Use with Caution’ have a lower correlation with their cognate mRNA measurements than protein measurements made with ‘Valid’ antibodies. The results in Fig suggest that it is not simply that these proteins display lower correlation with their cognate mRNAs as the same trend is not evident for mass spectrometry measurements.

To understand the variance in mRNA-protein correlation that can be associated with antibody validation status, we performed univariate linear regression. The antibody validation status could explain 5.5 - 18 % of the variation in the mRNA-protein correlation for the TCGA studies wherein the protein expressions are measured using RPPA (Fig S1). The average variance explained in the mRNA-protein correlation for the TCGA studies is ∼9%. However, the average variance explained in the mRNA-protein correlations for the CPTAC studies where proteins are measured using mass spectrometry is <1% (Fig S1). This suggests that the associated antibody validation status has a significant influence on mRNA-protein correlations when protein abundance is measured using RPPA but not when it is measured by MS.

### Antibody reliability does not reflect protein measurement reproducibility, protein abundance, or mRNA abundance

We have previously found that some proteins appear, across multiple studies, to be more reproducibly quantified by mass spectrometry than others (Upadhya & Ryan, 2022). We exploited this observation to develop an aggregated protein reproducibility rank for each protein that ranges from 0 to 1 (0 – not reproducible; 1 – highly reproducible). Using this score we found that proteins with more reproducible measurements tended to have higher observed mRNA-protein correlations across multiple mass spectrometry studies (Upadhya & Ryan, 2022).

To understand if proteins with less reliable antibodies are also less reproducibly measured by mass spectrometry, we compared the aggregated protein reproducibility ranks for the proteins with ‘Valid’ and ‘Use with Caution’ antibodies. We observed that there was no significant difference in the distribution of aggregated protein reproducibility ranks for the two groups of proteins (Fig 4A). This suggests that proteins with less reliable antibodies are not more irreproducible when measured using mass spectrometry and that therefore they are not inherently more challenging to quantify.

**Figure 4.**
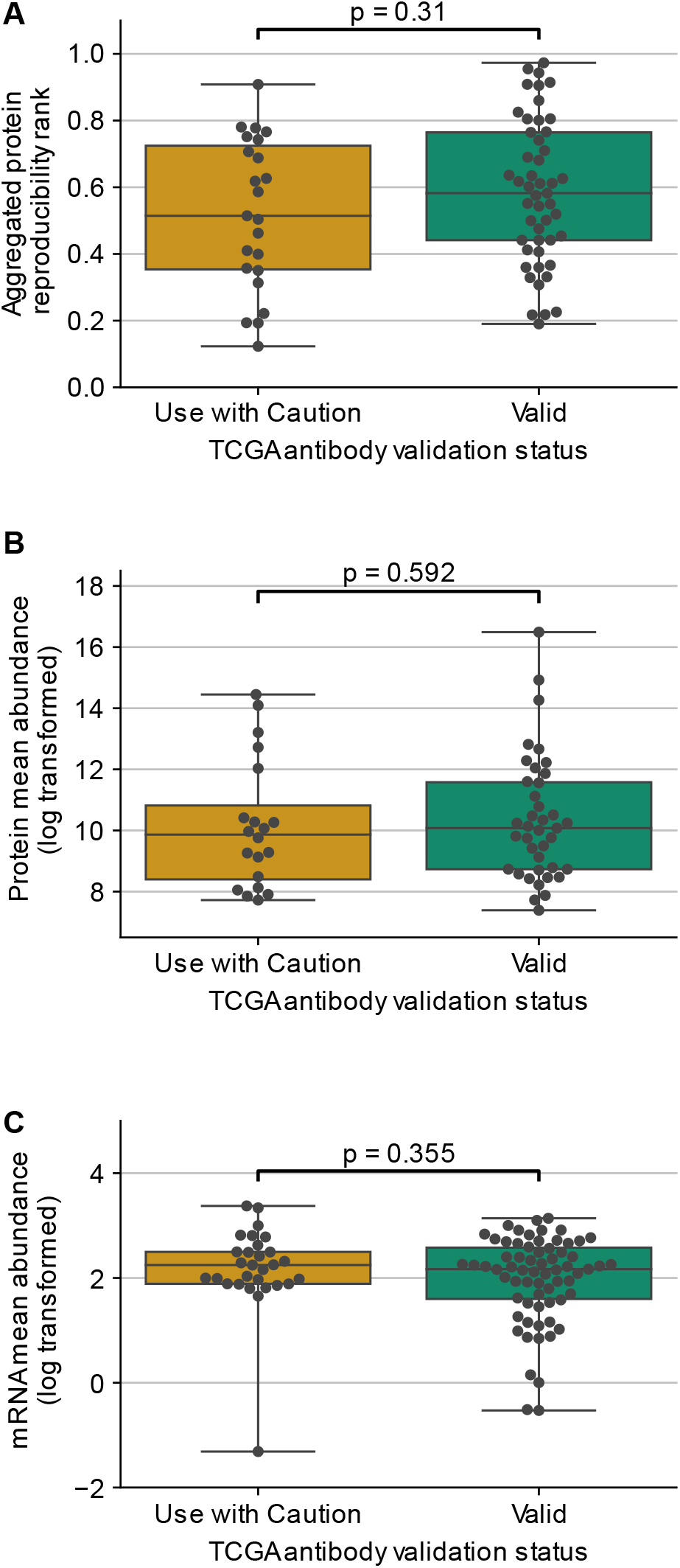
Antibody reliability does not reflect protein measurement reproducibility, protein abundance, or mRNA abundance. Boxplots showing the distribution of aggregated protein reproducibility ranks (A), protein abundances (B) and mRNA abundances (C) for proteins from TCGA studies classified based on their antibody validation status. For each box plot, the black central line represents the median, the top and bottom lines represent the first and third quartiles, and the whiskers extend to 1.5 times the interquartile range past the box. Each point on a box plot represents a protein.

We also previously found that protein abundance influences the reproducibility of protein measurements made using MS – more abundant proteins tend to be more reliably quantified by MS (Upadhya & Ryan, 2022). To test if antibodies marked as ‘Use with Caution’ preferentially target proteins with lower abundance, we compared the protein abundances of the targets of these antibodies to those targeted by ‘Valid’ antibodies. We obtained protein abundance measurements from the GTEx project (Jiang *et al*, 2020) and found that there was no significant difference in protein abundance between the proteins targeted by the two types of antibodies ‘Valid’ and ‘Use with Caution’ Fig The same trend was observed when we assessed mRNA abundances (Fig 4C).

Overall, these results suggest that the antibodies marked as ‘Use with Caution’ do not target proteins that are less abundant or that are less reproducibly quantified by mass spectrometry.

### Validation of findings using the Cancer Cell Line Encyclopedia

Thus far we have focussed our analysis on tumour samples profiled by either the TCGA or CPTAC. We have used the same, or similar, cancer types when analysing mRNA-protein correlations in both studies. However, a limitation of our analysis is that we have not been able to use the same sets of samples for all analyses – this is because many of the CPTAC MS studies do not have associated RPPA data available while the majority of TCGA samples profiled by RPPA do not have corresponding MS profiles. Consequently we have assessed mRNA-protein correlations for RPPA and MS protein quantification using different sets of samples.

The Cancer Cell Line Encyclopedia, and associated molecular characterisation efforts, presents an opportunity to validate our findings in an orthogonal dataset where the same cell lines have been profiled using RPPA, MS, and RNASeq. 359 cancer cell lines have measurements for all three data types available through the DepMap portal (Ghandi *et al*, 2019; Nusinow *et al*, 2020). The RPPA data used a panel of 155 antibodies to quantify individual, non-phosphorylated proteins where 79 antibodies overlapped with the TCGA antibodies. We identified that ∼ of the CCL antibodies were marked as ‘Use with Caution’ Fig A while the remainder were marked as ‘Valid’. As with our analysis of the TC A data, we found that proteins quantified using antibodies marked as ‘Use with Caution’ have lower mRNA-protein correlation than those measured with ‘Valid’ antibodies (Fig .5B). Furthermore, as in the TCGA, when the same proteins were quantified using mass spectrometry, there was no significant difference in the mRNA-protein correlation between the two groups (Fig 5C). Similar to the TCGA antibodies, the proteins in the CCLE study with ‘Valid’ and ‘Use with Caution’ antibodies had no significant difference in protein reproducibility, protein or mRNA abundances (Fig S2).

**Figure 5.**
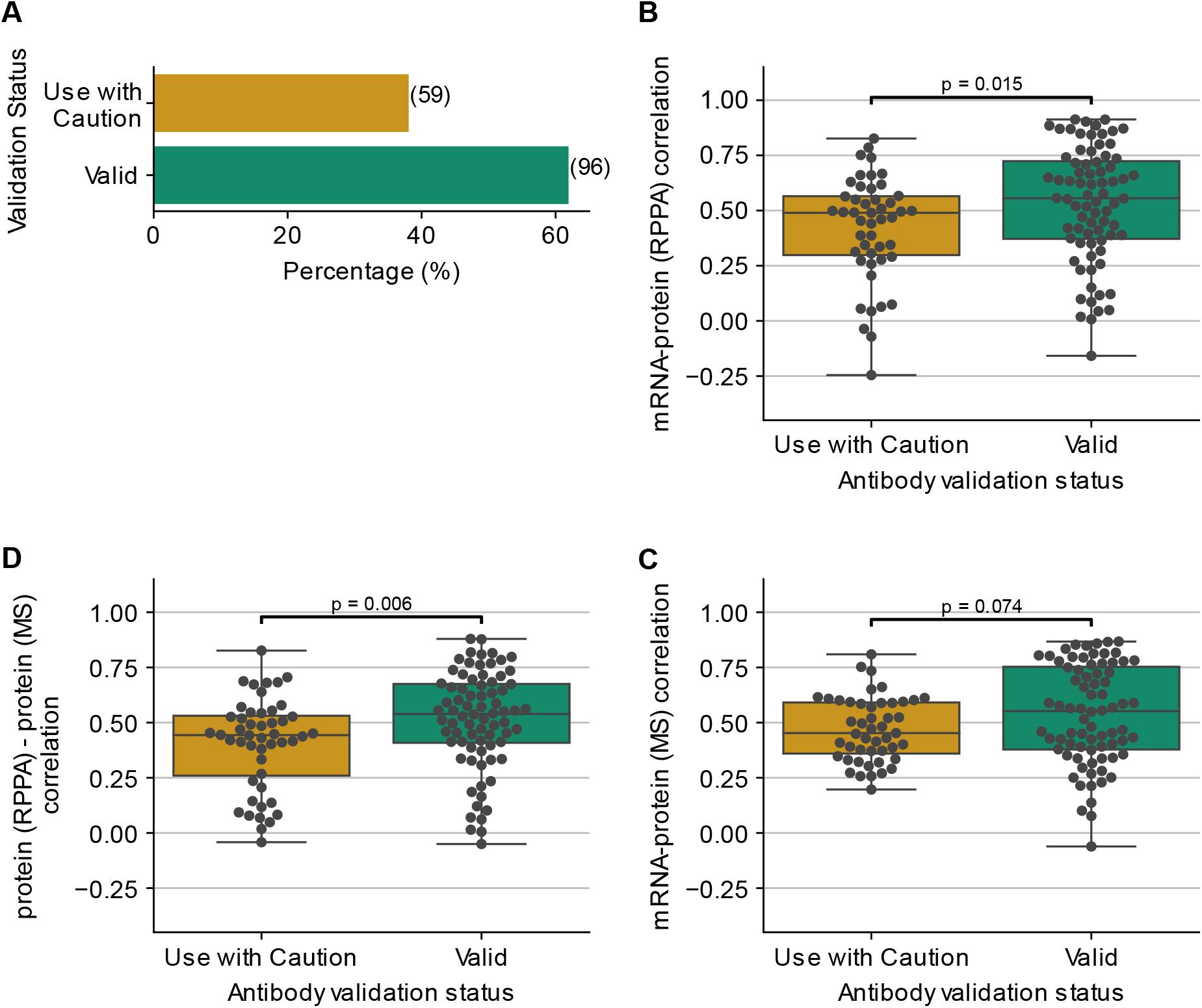
Validating the influence of antibody validation status on mRNA-protein correlation on CCLE data. (A) Bar chart showing the percentage of antibodies used to quantify proteins in the cancer cell lines based on the antibody validation status. The actual number of antibodies in each class is shown in brackets. (B) Boxplots showing the distribution of mRNA-protein correlation for proteins quantified using antibodies labelled as ‘Valid’ or ‘Use with Caution’. C oxplots showing the distribution of mRNA-protein correlations wherein proteins are classified based on their antibody validation status and quantified using mass spectrometry. (D) Boxplots showing the distribution of protein (RPPA) - protein (mass spectrometry) correlations based on their antibody validation status. For each box plot, the black central line represents the median, the top and bottom lines represent the first and third quartiles, and the whiskers extend to 1.5 times the interquartile range past the box. Each point on a box plot represents a protein.

The availability of protein abundance measurements from two different techniques (RPPA and MS) over the same cancer cell lines enables us to directly compare the protein abundance measurements obtained from the two (RPPA and MS) techniques. We calculated the correlation between protein abundance measurements obtained from RPPA and MS for each protein and grouped the proteins based on their antibody validation status. Proteins with valid antibodies had a significantly higher median protein (RPPA) – protein correlation compared to the proteins labelled ‘Use with Caution’ Fig D p-value = 0.006, Mann-Whitney U test, two-sided). This suggests that protein abundance measurements made with ‘Use with Caution’ antibodies show less concordance with measurements of the same proteins. Overall this is consistent with these measurements being less reliable and consequently contributing to the lower observed mRNA-protein correlations.

## Discussion

In previous work we have demonstrated that some proteins appear to be more reproducibly quantified by mass spectrometry than others, and that more reproducibly quantified proteins tend to have higher observed mRNA-protein correlations (Upadhya & Ryan, 2022). Here, we demonstrate that the mRNA-protein correlations observed using RPPA quantification measurements are significantly influenced by the reliability of the antibodies used. We find that proteins that are quantified by RPPA using more reliable antibodies tend to have higher observed mRNA-protein correlations. This is not true when the same proteins are quantified by mass spectrometry, suggesting that the cause is likely the antibodies themselves rather than the measured proteins somehow being less correlated with their associated mRNAs. Further support for this comes from the observation that RPPA protein measurements are more highly correlated with MS measurements of the same proteins when the RPPA measurements are made using ‘Valid’ antibodies.

Although collectively often referred to as high quality antibodies, almost one third of the antibodies used by the TC A are labelled as ‘Use with Caution’. The list of antibodies, and their validation status, are made available in the supplementary materials of relevant publications (Li *et al*, 2013; Akbani *et al*, 2014b) and associated online resources. However, the full set of R A measurements, including those made with antibodies marked as ‘Use with Caution’, are still often used for systematic analyses (Zhang *et al*, 2015, 2017; Koplev *et al*, 2018; Chen *et al*, 2019). Furthermore the measurements made with these antibodies are made available through widely used web resources that facilitate browsing of TCGA data without any indication that these measurements are less reliable (Cerami *et al*, 2012; Gao *et al*, 2013; Vasaikar *et al*, 2018). Our results suggest that the measurements from these antibodies should be used with caution in systematic analyses and that they should also be flagged as ‘Use with Caution’ in relevant web resources.

We note that the measurements made with antibodies labelled ‘Use with Caution’ may still be reproducible but that they provide a less accurate quantification of the target protein and hence display lower mRNA-protein correlations. It may be the case that they measure the joint abundance of multiple proteins and that this aggregate measurement may be reproducible, but still not reliable.

Here, we have focussed on the analysis of RPPA profiles of tumour samples and cancer cell lines, but RPPA profiling is also used in other contexts. For instance, they have been used to systematically profile the responses of cell lines to systematic perturbations (Korkut *et al*, 2015; Keenan *et al*, 2018) and also to understand profiles of patients with diseases other than cancer (Napierala *et al*, 2021). The results of such studies are also likely to be impacted by the use of antibodies of varying quality.

The issue of antibody reliability is a general challenge for biological studies, not just those that make use of RPPA (Baker, 2015; Goodman, 2018). By performing systematic evaluations of all antibodies used, the TCGA RPPA studies make use of antibodies that are likely of significantly higher quality than average. Even those antibodies marked ‘Used with Caution’ are still deemed of sufficient quality for inclusion in assays and are likely to be of higher quality than randomly selected antibodies. Therefore the trends observed in our analysis are likely to be a lower bound of the potential impact of unreliable antibodies on protein abundance measurements.

## Materials and Methods

### Data collection

The transcriptomic and RPPA data for the TCGA Pan-Cancer Atlas studies (breast, ovarian, colorectal, endometrial, kidney and brain) were downloaded from USC Xena browser (Goldman *et al*, 2020). For the analyses using CPTAC studies, the mRNA-protein correlations were obtained from the supplementary table of our previous publication (Upadhya & Ryan, 2022). The transcriptomic profiles in all studies were quantified using RNA-Seq. The protein expression profiles of TCGA studies were quantified using RPPA whereas the CPTAC studies had proteomic profiles quantified using mass spectrometry. For the CCLE study, the mRNA expression (Ghandi *et al*, 2019), mass spectrometry based protein expression (Nusinow *et al*, 2020) and RPPA protein expression profiles (Ghandi *et al*, 2019) were downloaded from the cancer dependency map portal (https://depmap.org/portal/ccle/).

### Pre-processing protein and transcript expression

The protein expression profiles quantified using RPPA contained missing values for a small number of proteins. Within each study we restricted our analyses to proteins that were measured in at least 80% of samples. Some antibodies measure the protein products of multiple genes – e.g. the AKT antibody (Akt) measures the total protein abundance from the protein products of the genes *AKT1, AKT2, and AKT3*. These antibodies that target multiple proteins were excluded from our analysis. Some antibodies are used to measure the abundance of specific phosphoproteins (e.g. 4E-BP1_pS65 measures a specific EIF4EBP1 phosphoprotein) rather than total protein abundances. These were also excluded from our analyses. The criteria for retaining the transcript for the analyses was the same as that for protein – measured (non zero values) in at least 80% of samples.

### Antibody annotation

The antibodies for the TCGA Pan-Cancer Atlas studies were obtained from the supplementary data of (Li *et al*, 2013). There were 187 antibodies marked as ‘Use with Caution’, ‘Valid’ 11 and ‘Under valuation’ 2. We restricted our analyses to antibodies that (i) mapped to no more than one protein (ii) did not map to phosphoproteins iii were marked as ‘Valid’ or ‘Use with Caution’ only. ased on the first two criteria, we had 132 antibodies (78 labelled as ‘Valid’, 30 labelled as ‘Use with Caution’ and 2 labelled as ‘Under valuation’. For the antibodies marked as ‘Under valuation’, we obtained the current validation status for 7 additional antibodies from the validated antibody standard list from MD Anderson Cancer Center website (https://www.mdanderson.org/research/research-resources/core-facilities/functional-proteomics-rppa-core/antibody-information-and-protocols.html). The incorrect or the uncommon gene names in the standard list of antibodies from the aforementioned webpage were corrected as indicated on the website.

Overall, we were able to analyse 114 proteins with known antibody validation status for the TCGA Pan-Cancer Atlas studies (Table S1). The antibodies that are not used for the analyses in this study have been included in Table S2.

There were 214 antibodies for the CCLE cancer cell lines study that were obtained from the supplementary data of (Ghandi *et al*, 2019). Of these 214 antibodies, 139 antibodies were marked as ‘Valid’ and ‘7 ‘ marked as ‘Use with Caution’. ased on the criteria for our analyses, the number of antibodies reduced to 155 antibodies for the cancer cell lines.

### Computation of correlation coefficient

Correlation between mRNA-protein(RPPA) and protein(RPPA)-protein(MS) was computed using the Spearman rank correlation. For each protein in each study, samples with missing values were ignored when computing the correlation.

### Assessing the relationship between antibody validation status and mRNA-protein correlation (Figure S1)

To understand the variance in mRNA-protein correlation explained by antibody validation status we used linear models given by the equation –

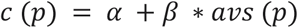

where *c* (*p*) is the mRNA-protein (protein expression measured through either RPPA or MS) correlation for each protein, *avs* (*p*) is the antibody validation status for each protein represented as 1 if the antibody is ‘Valid’ and 0 if the antibody is marked as ‘Use with Caution’. The coefficients *α* and *β* are estimated using the ordinary least squares regression method. All linear regressions were carried out using the statsmodel package in Python.

### Statistical Analysis

All statistical analyses were carried out using Python 3.11.0, Pandas 1.5.2 (Mc Kinney, 2011), numpy 1.24.0 (Harris *et al*, 2020), scipy 1.9.3 (Virtanen *et al*, 2020) and statsmodels 0.13.5 (Seabold & Perktold, 2010). The figures were created with Matplotlib 3.6.2 (Hunter, 2007) and Seaborn 0.12.1 (Waskom, 2021).

## Data Availability

This paper analysed existing, publicly available data. All original code has been deposited at Github (https://github.com/cancergenetics/antibody_quality_limitations.git).

## Acknowledgements

S.R.U was funded through the School of Computer Science, University College Dublin and C.J.R was funded by the Irish Research Council Laureate Awards 2017/2018. We thank members of the Ryan lab for careful reading of the manuscript and helpful feedback.

## Author contributions

Conceptualization, S.R.U and C.J.R; Methodology, S.R.U and C.J.R; Formal analysis, S.R.U; Data curation, S.R.U; Writing – original draft, S.R.U and C.J.R; Writing – review and editing, C.J.R; Visualization, S.R.U; Supervision, C.J.R ; Funding acquisition, C.J.R

## Conflict of interest

The authors declare no competing interests.

## Supplementary Files

**Table S1. Antibodies used for analysing TCGA Pan-Cancer RPPA studies**

The list of antibodies used to analyse the TCGA Pan-Cancer RPPA data in this study.

**Table S2. Antibodies that are not used for our analyses**

The list of antibodies that are not being used for the analyses in this study and the associated reasons

## Supplementary Figures

**Figure S1.**
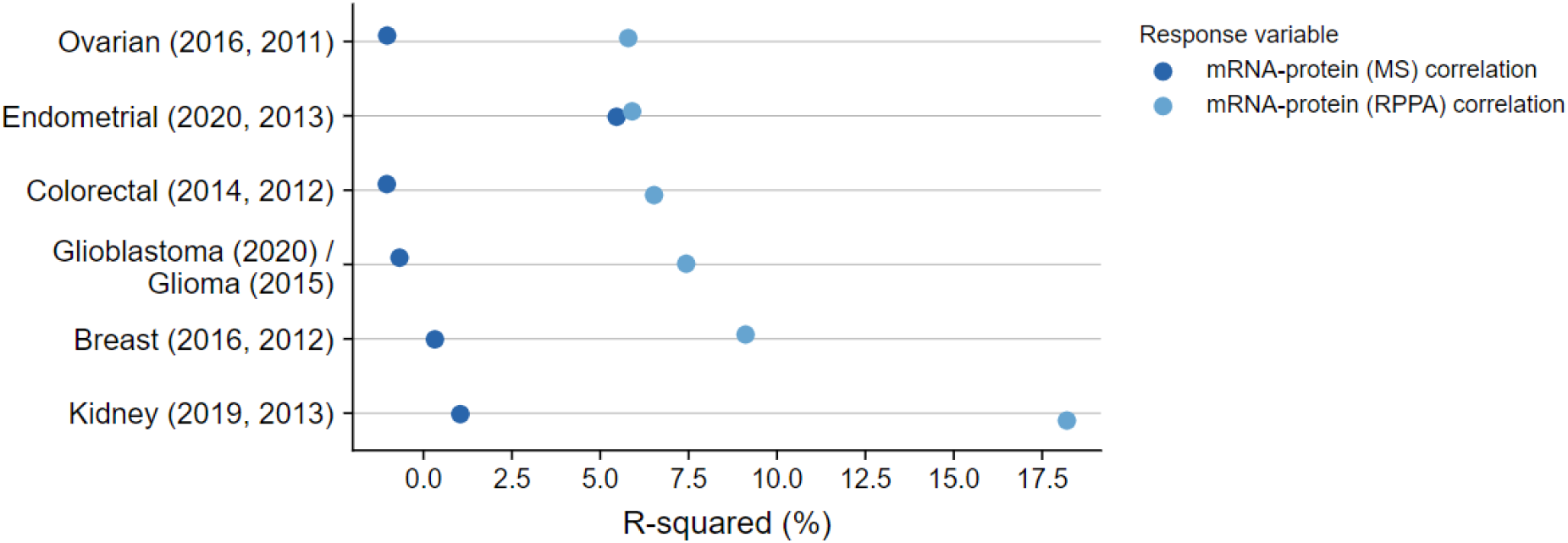
Validating the influence of antibody validation status on mRNA-protein correlation using univariate linear regression. Dot plot comparing the R-squared obtained by regressing the antibody validation status from mRNA-protein (RPPA and MS) correlation for each pair of studies (CPTAC, TCGA).

**Figure S2.**
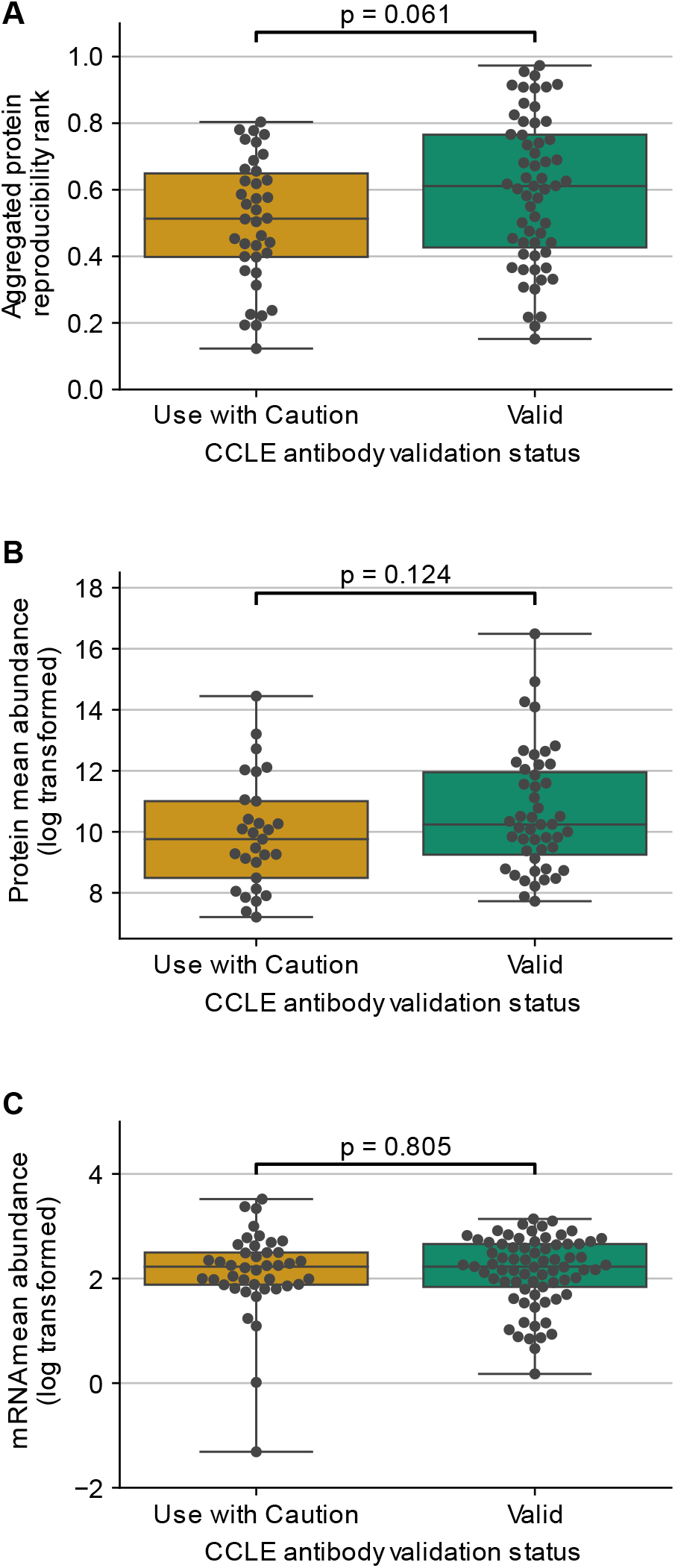
Validating that antibody reliability does not reflect protein measurement reproducibility, protein abundance, or mRNA abundance on CCLE data. Boxplots showing the distribution of aggregated protein reproducibility ranks (A), protein abundances (B) and mRNA abundances (C) for proteins from cancer cell lines classified based on their antibody validation status. For each box plot, the black central line represents the median, the top and bottom lines represent the first and third quartiles, and the whiskers extend to 1.5 times the interquartile range past the box. Each point on a box plot represents a protein.

